# BayesDeBulk: A Flexible Bayesian Algorithm for the Deconvolution of Bulk Tumor Data

**DOI:** 10.1101/2021.06.25.449763

**Authors:** Francesca Petralia, Azra Krek, Anna P. Calinawan, Daniel Charytonowicz, Robert Sebra, Song Feng, Sara Gosline, Pietro Pugliese, Amanda G. Paulovich, Jacob J. Kennedy, Michele Ceccarelli, Pei Wang

## Abstract

To understand immune activation and evasion mechanisms in cancer, one crucial step is to characterize the composition of immune and stromal cells in the tumor microenvironment (TME). Deconvolution analysis based on bulk transcriptomic data has been used to estimate cell composition in TME. However, these algorithms are sub-optimal for proteomic data, which has hindered research in the rapidly growing field of proteogenomics. Moreover, with the increasing prevalence of multi-omics studies, there is an opportunity to enhance deconvolution analysis by utilizing paired proteomic and transcriptomic profiles of the same tissue samples. To bridge these gaps, we propose BayesDeBulk, a new method for estimating the immune/stromal cell composition based on bulk proteomic and gene expression data. BayesDeBulk utilizes the information of known cell-type-specific markers without requiring their absolute abundance levels as prior knowledge. We compared BayesDeBulk with existing tools on synthetic and real data examples, demonstrating its superior performance and versatility.

**Availability:** Software available at http://www.BayesDeBulk.com/

**Contact:** For any information, please contact

## 1. Introduction

Tumor tissues comprise various cell types, including immune and stromal cells. Quantifying the proportion of different cell types in the tumor microenvironment (TME) is crucial for understanding immune activation and evasion mechanisms as well as their roles in tumor progression. Over the past decade, many algorithms have been proposed to estimate TME cell compositions based on bulk transcriptomics data. However, these deconvolution algorithms are often suboptimal for analyzing other data types, such as proteomic profiles, as different omics data have distinct features. Moreover, in recent cancer studies [1, 2, 3, 4], it became common to collect multi-omic data, including both transcriptomic and proteomic data, for each biological sample. Then, it is ideal to fully leverage all available data in order to better estimate the immune and stromal composition in the TME. To bridge these gaps, in this work, we propose a new method called BayesDeBulk for estimating the immune/stromal cell composition based on bulk proteomic or multi-omics profiles.

Existing deconvolution methods, which were mainly developed for transcriptomic data, can be categorized into three major types: reference-based, semi-reference-based, and reference-free methods. The reference-based methods, such as CIBERSORT and CIBERSORTx [5, 6, 7, 8, 9, 10], require prior information of gene expression levels in individual cell types. However, issues may arise when the gene expressions of certain cell types in different organs or tumors do not align with those in the reference data. These issues are especially prevalent when analyzing mass-spectrometry based proteomics data, in which non-ignorable batch effects exist across different experiments. To address this lack of flexibility, semi-reference-based approaches [11, 12], including Bayesian algorithms [13, 14] have been developed for deconvolution analysis. The latter utilizes prior distributions based on single cell RNA (scRNA) sequencing profiles. However, without proper constraints on the parameter space, the resulting posterior distributions of gene expressions may deviate considerably from the prior distribution posing identifiability issues of the estimated components. Furthermore, utilizing such models can be challenging in practice when suitable scRNA data is not available.

Among the reference-free methods, xCell [15] is one of the most commonly used. xCell considers a list of markers expressed in each cell type (cell-type specific markers) as prior information and returns enrichment scores reflecting the amount of different cell types in the TME. While xCell is more flexible than other reference-or semi-reference-based deconvolution algorithms, it only estimates cell type fractions without providing any information on cell type-specific gene expression levels, which is often needed to characterize the status of individual cell types for downstream analysis. Several other reference-free algorithms aim to infer both cell-type fractions and cell type-specific (mean) marker expressions, such as Plier, Pathway-level Analysis of Transcriptomic Signatures (PATS), TOAST, and RefFreeEWAS [16, 17, 18, 19, 20]. These algorithms use factor analysis or similar strategies without relying on reference cell-type gene expression profiles. However, the results can be difficult to interpret since it is often challenging to understand which cell type(s) are represented by each inferred factor. Wang et al [20] proposed DemixT which can overcome this identifiability problem leveraging markers expressed by each cell type. However, this algorithm can be applied to capture only three cell-types (i.e., stromal, immune and tumor cells) and, therefore, it fails to characterize higher resolution cell-type composition. Mao et al [16] proposed Plier, a hierarchical factor model that utilizes information of cell-type specific markers, but the lack of a one-to-one mapping between inferred factors and cell types poses considerable interpretability issues of the estimated factors. Recently, Tang et al [21] introduced an algorithm based on non-negative matrix factorization, which solves the identifiability problem by employing a penalized regression model to shrink towards zero the gene expression estimates of markers expected to be less expressed in a particular cell type. However, in practice, it is often difficult to accurately label genes with three categories (not expressed, expressed, and highly expressed) for each cell type, which are required by the algorithm as input. In addition, all of the above methods cannot be used to simultaneously deconvolve multi-omics data derived from the same set of samples.

In this work, we propose BayesDeBulk, a novel reference-free method for bulk deconvolution analysis. The method utilizes a list of cell-type specific markers and imposes a Repulsive prior distribution on the mean of these markers’ expression to ensure their upregulation in the corresponding cell-type components. Repulsive prior is a class of priors introduced by Petralia et al [22] that ensures the interpretability and identifiability of mixture models, and has been recently extended to different applications [23, 24, 25, 26]. Furthermore, BayesDeBulk models the cell-type fraction parameters using a spike-and-slab prior [27], which induces sparsity to screen away cell types which are not present in the tumor microenvironment from the model. BayesDeBulk is particularly suitable for deconvolution analysis based on proteomics data, because appropriate reference data for cell type-specific protein signatures are usually not available. Moreover, unlike existing reference-free methods, the Repulsive prior in BayesDeBulk uniquely determines the mapping between different components and cell types. Additionally, the mixture model of BayesDeBulk supports the joint modeling of multi-omic data, such as proteomic and transcriptomic data, measured for the same set of samples. Superior performance of BayesDeBulk over existing deconvolution methods such as Cibersort [5], CibersortX [9], xCell [15], Epic [6], Plier [16], and MCP-counter [10] is shown on both synthetic and real datasets, highlighting its versatility and value.

## 2. Results

### 2.1. BayesDeBulk

BayesDeBulk can be applied to individual omics dataset (e.g., transcriptomic or proteomic data) or the integration of omics datasets. For the sake of simplicity, in the below narrative, we consider protein expression data as an example to describe the method.

Protein expression data of bulk tissue can be viewed as the weighted average of the protein expression profiles of different cell types in the tissue. Specifically, we model the expression of protein *j* for the *i*-th sample as a Gaussian distributed random variable with mean parameter being the linear combination of the expression of protein *j* in different cell types as follows:

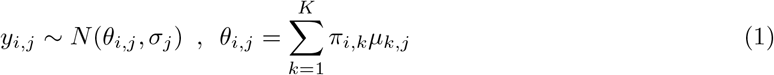

where *K* is the total number of cell types in the tissue, *π*_*i,k*_ is the fraction of the *k*-th cell type in the *i*-th sample, *µ*_*k,j*_ is the expression of protein *j* in the *k*-th cell type and *σ*_*j*_ is the variance of the *j*-th protein. For reference-based methods, *µ*_*k,j*_ are generally treated as known parameters with values derived from reference data of purified cells [5, 7, 8]. For reference-free approaches, *µ*_*k,j*_ shall be estimated from the data. However, without proper constraints on the parameter space, model in (1) is often not identifiable. To overcome this issue, we propose to use a Bayesian approach with a Repulsive prior specified on the mean parameters {*µ*_*k,j*_} [22]. Let us assume that for the *k*-th cell type, there is a set of markers whose expressions are upregulated in the *k*-th cell type compared to the *s*-th cell-type (see Figure 1-A). The index of this set can be annotated as:

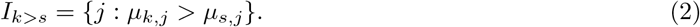

We use a flexible Repulsive prior [22] to ensure that markers in set *I*_*k>s*_ will have a “larger” mean in the *k*-th cell type compared to the *s*-th cell-type. Denote 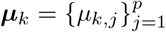 as the mean expression vector of *p* markers in the *k*-th cell type. Then, (***µ***_1_, …, ***µ***_*K*_) is jointly modeled through the following multivariate prior distribution:

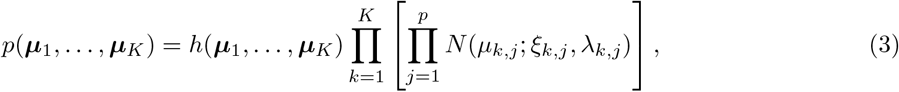

where *ξ*_*k,j*_ and *λ*_*k,j*_ are the mean and the variance parameters of the Gaussian prior distribution, and *h*(·) is a Repulsive function defined as

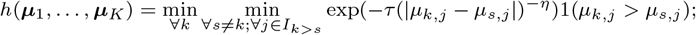

with 1(*A*) being an indicator function equal to 1 if *A* is satisfied and 0 otherwise, *τ* being a positive parameter, and *η* a positive integer controlling the rate at which the Repulsive function approaches zero.

**Figure 1:**
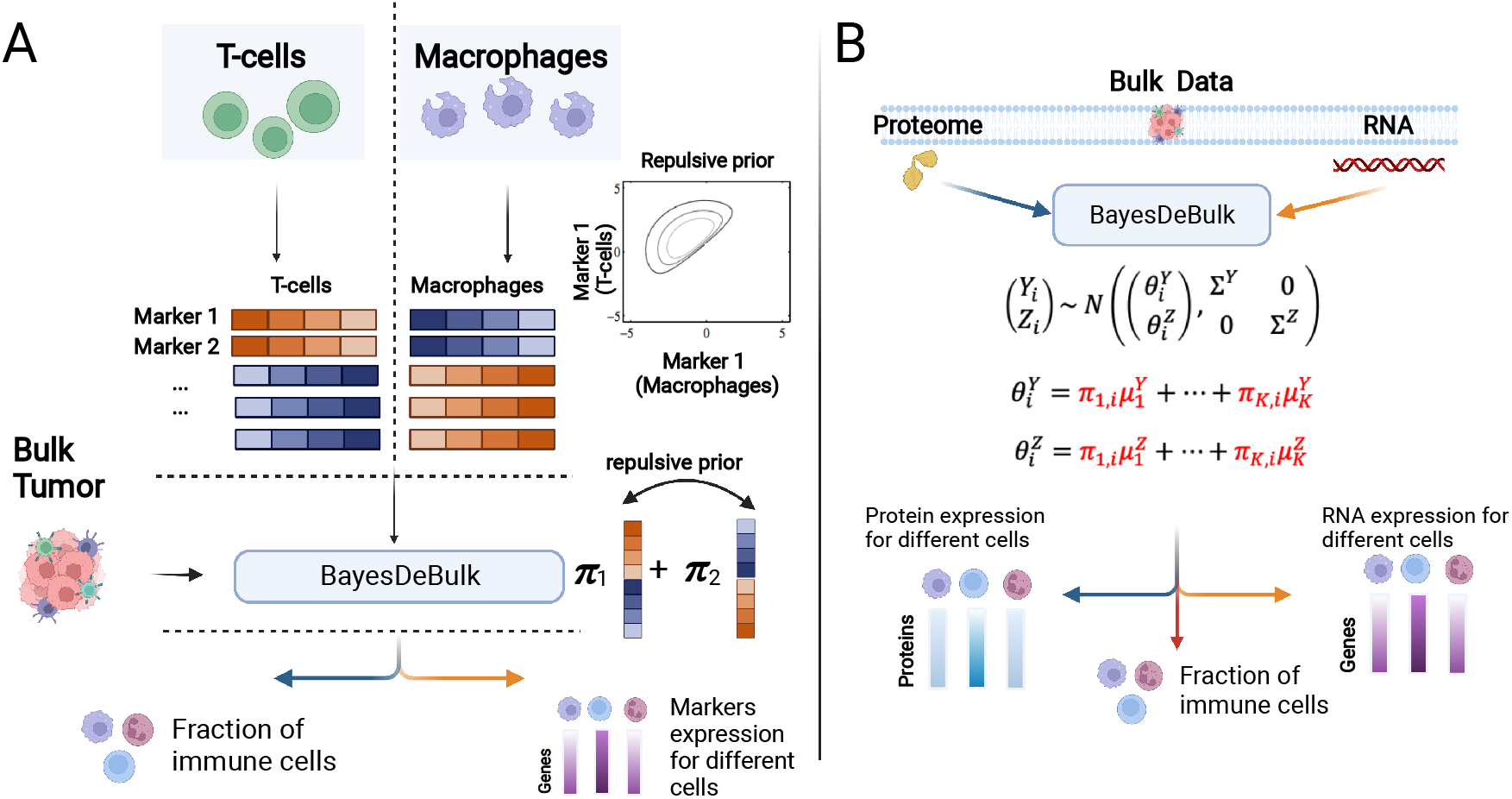
Algorithm Schematic. (A) Bulk data is modeled as a linear combination of marker expression in different cell types. Given a list of markers expressed in each cell type, a Repulsive prior is placed on the mean of marker expression in different cell types to ensure that cell type specific markers are upregulated in a particular component. (B) Multi-omic framework to estimate cell type fractions integrating proteomic and RNAseq data. Given a list of cell-type specific markers, the algorithm returns the estimated protein/RNA expression for different cell types and cell-type fractions for different samples.

The *h*(·) function is an extension of the Repulsive function introduced by Petralia et al [22]. According to this function, markers contained in set *I*_*k>s*_ will have larger mean values in the *k*-th cell type than in the *s*-th cell type. Specifically, it takes value 0 if the mean parameter of any marker from *I*_*k>s*_ for the *k*-th cell type are not higher compared to that of the *s*-th cell type. Otherwise, it takes a positive value, and increases with the absolute distance between mean parameters of the two cell types. Note that, in (3), Repulsive priors are imposed only on markers belonging to the set *U*_*k,s*_*I*_*k>s*_, while Gaussian prior distributions are used for other markers. This approach ensures model identifiability and reduces the computational burden. Prior knowledge from existing databases and/or single-cell data can be used to specify *I*_*k>s*_. The Repulsive prior also allows users to incorporate differential expressed protein list for each pair of cell-types. This is helpful since it is often problematic to find markers upregulated in a particular cell type compared to all other cell types. For instance, the set of markers upregulated in Naive CD4 T Cells compared to Myeloid cells could be very different from the set of markers upregulated in Naive CD4 T Cells compared to Memory CD4 T Cells. By enabling users to specify differential markers for each pair of cell types, BayesDeBulk effectively leverages all relevant prior information.

A standard choice for the parameters of the Gaussian prior is *ξ*_*k,j*_ = 0 and *λ*_*k,j*_ = 1. Alternatively, these parameters can be chosen based on prior knowledge from existing databases and single-cell datasets. While each *π*_*i,k*_ is defined on the unit interval [0, 1], for the ease of computation, BayesDeBulk does not require the sum of 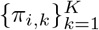 being equal to 1. For the prior specification for *π*_*i,k*_, BayesDeBulk uses a spike-and-slab prior [27] defined on the unit interval (Star Method).

BayesDeBulk can be utilized to perform the deconvolution via matched gene and protein expression data (Figure 1-B). In this case, each data type can be modeled via separate mixture models sharing the same set of cell type fraction parameters (Star Method). It is important to notice that the proposed flexible framework allows the user to integrate both proteomic and RNAseq data to infer cell type fractions for samples for which the two data types are measured; and a single-omic deconvolution for sample with one data type measured.

### 2.2. Simulated Data

#### 2.2.1. Evaluation of parameters estimates based on simulated data

The performance of BayesDeBulk in estimating cell-type fractions and the marker expression in different cell types was evaluated based on extensive synthetic data. Pseudo bulk data was generated from a Gaussian model as explained in details in the Star Method (Figure S1). BayesDeBulk was compared with Cibersort [5], Plier [16], xCell [15] and EPIC [6] based on different simulation scenarios with varying numbers of cell types *K*, markers *p* and sample size *n*; i.e., (*K, p, n*) = (10, 200, 50), (*K, p, n*) = (20, 400, 50), (*K, p, n*) = (10, 200, 100) and (*K, p, n*) = (20, 400, 100), and variance levels *ν* and *σ*. For each synthetic scenario, 30 replicate datasets were generated and the performance of different models was evaluated based on two metrics: Pearson’s correlation and mean squared error (MSE) between estimated fractions and true fractions. For each replicate, BayesDeBulk was estimated considering 10, 000 Marcov Chain Monte Carlo (MCMC) iterations; with the estimated fractions being the mean across iterations after discarding a burn-in of 1, 000. Different methods were implemented (i) assuming that all cell-type specific markers are known a priori and (ii) only 50% of cell-type specific markers are known a priori. This second scenario is more representative of real world applications, where only a proportion of cell-type specific markers is usually known. Cibersort and EPIC require as input a signature matrix containing the expression of different markers for different cell types. In order to make a fair comparison, a perturbed version of the original signature matrix was considered as input (Star Method).

Figure 2 shows the performance of all the models in estimating cell-type fractions for different synthetic data scenarios considering a sample size *n* = 100 under the assumption that only 50% of cell-type specific markers is known. As shown, BayesDeBulk resulted in the highest Pearson’s correlation and lowest MSE between estimated and true cell-type fractions for different synthetic data scenarios. A median correlation above 0.90 was observed for BayesDeBulk for all simulation scenarios involving *K* = 10 components. Among other deconvolution algorithms, Plier resulted in the best performance in terms of Pearson’s correlation with a median correlation greater than 0.85 for all synthetic scenarios involving *K* = 10 cell types, although leading to the worst MSE. In addition, given the identifiability issue affecting factor models, for many synthetic scenarios, Plier was not able to map factors to a particular cell type (Star Method). This identifiability issue is expected to be more pronounced in real-world applications, posing considerable challenges to Plier’s implementation. As expected, the performance of all models decreased as more cell types were considered (K=20). Given their ability to estimate the expression of different markers for multiple cell types, BayesDeBulk and Plier were also compared in this regard. Figure 2 shows the performance of BayesDeBulk and Plier in estimating the mean of marker expression for different components. As expected, higher noise levels resulted in lower performance in terms of both correlation and MSE. The median Pearson’s correlation between estimated and true values across replicates was above 0.80 for the simulations involving 10 cell types. Although the median correlation decreased substantially when the number of components increased to *K* = 20, it remained above 0.50 for different simulation scenarios. Additional synthetic data scenarios based on lower sample size *n* = 50 and different degree of prior knowledge can be found in Supplementary Figures 2, 3 and 4. Overall, we observe that the performance of BayesDeBulk is not affected by different degree of prior knowledge contrary to Cibersort, Epic and xCell.

**Figure 2:**
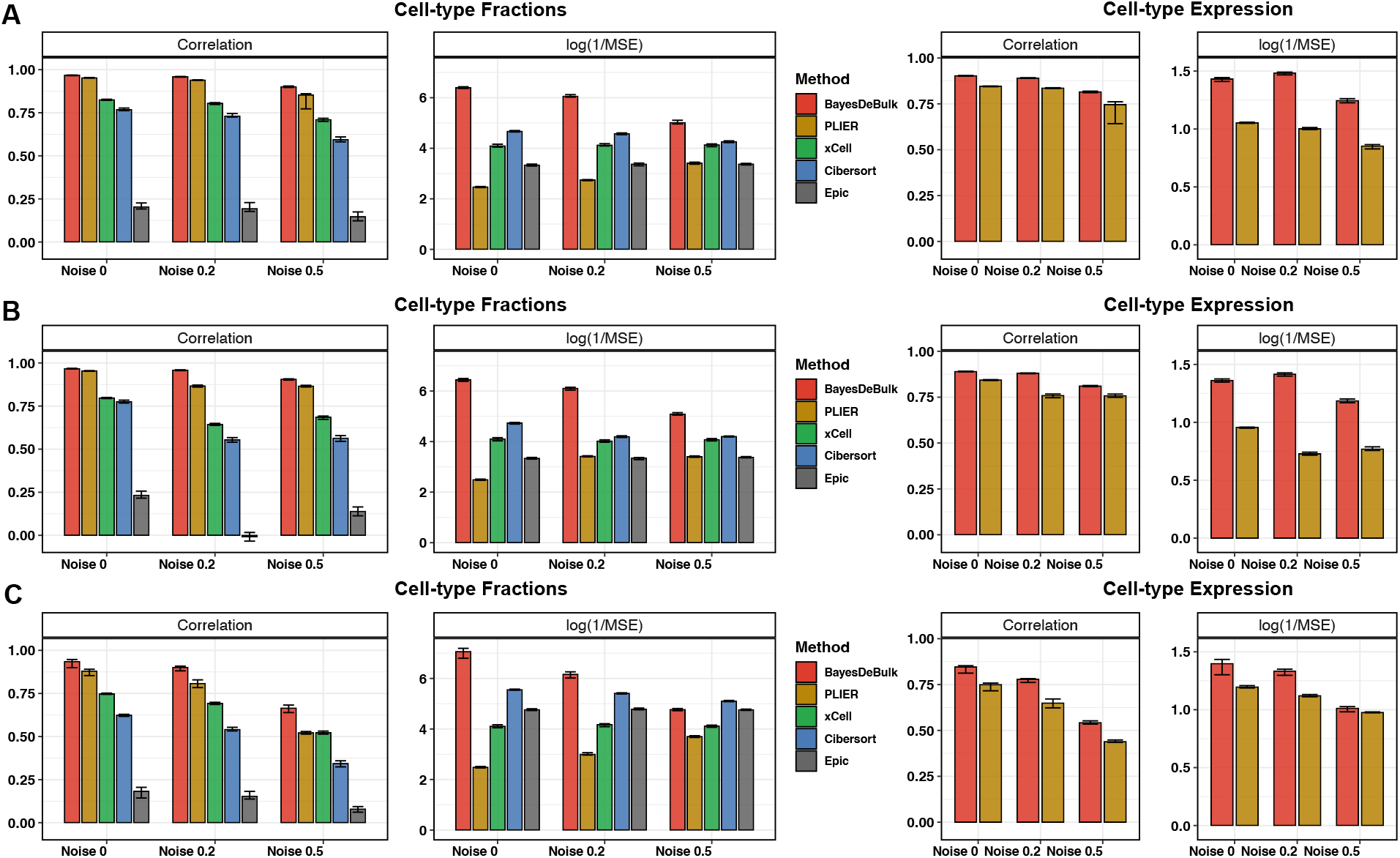
Pearson’s correlation and mean squared error (MSE) between estimated values and true values over 30replicates for BayesDeBulk (red), Cibersort (blue), xCell (green), Epic (gray) and Plier (gold). Barplots correspond to the median across different replicates while error bars to the interquartile range. For each simulation scenario, we report the correlation and MSE between the estimated cell-type fractions and the true values (left-hand panel) for all five algorithms, and between the estimated cell-type expression and true values (right-hand panel) for BayesDeBulk and Plier. Results are based on data simulated for (A) *K* = 10 and *σ* = 0.5; (B) *K* = 10 and *σ* = 1; (C) *K* = 20 and *σ* = 0.5 for different level of measurement errors *ν* (noise).

#### 2.2.2. Validation based on proteomic and transcriptomic pseudo-bulk data from purified cells

In this section, we compare different algorithms based on mixture of gene and protein expression datasets from purified cells. For this experiment, we considered data from [28] which contains transcriptomic profile of *K* = 6 immune cell types such as Neutrophil, Natural Killers, B cells, CD4 T cells, CD8 T cells and Monocytes. Considering the transcriptomic profiles of these immune cells, we obtained pseudo-bulk RNA data (Star Method). Then, we considered data from Rieckmann et al [29] including proteomic profiles for the same set of immune cells. Mixed proteomic data was generated in a similar fashion as the transcriptomic profile, considering the same set of mixture proportions. BayesDeBulk was compared with Cibersort [5], Epic [6], Plier [16], xCell [15] and MCP-counter [10] in estimating immune cell-type fractions. Considering a sample size of *n* = 50, 30 replicate datasets were generated and the performance of the models were evaluated based on the correlation between estimated and true fractions (Star Method). All algorithms were implemented using proteomic and transcriptomic data. For BayesDeBulk, a multi-omic based learning was also implemented.

Figure 3 shows the Spearman’s correlation and mean squared error (MSE) between true and estimated cell-type fractions for different algorithms. Overall, BayesDeBulk resulted in the highest correlation and lowest MSE. For some cell types such as Neutrophils, the multi-omic based deconvolution was able to slightly outperform single-omic based deconvolutions revealing the advantage of a multi-omic based learning. Although Plier was the algorithm with the highest correlation among competitors, it resulted in the highest MSE. Similarly to other data examples, for a large number of replicates (*∼* 50%), Plier could not map estimated factors to each one of the 6 immune cell types. Only replicates for which this mapping was possible were considered to produce summary statistics in Figure 3. As previously highlighted, this identifiability problem is an issue when applying Plier to real-world data.

**Figure 3:**
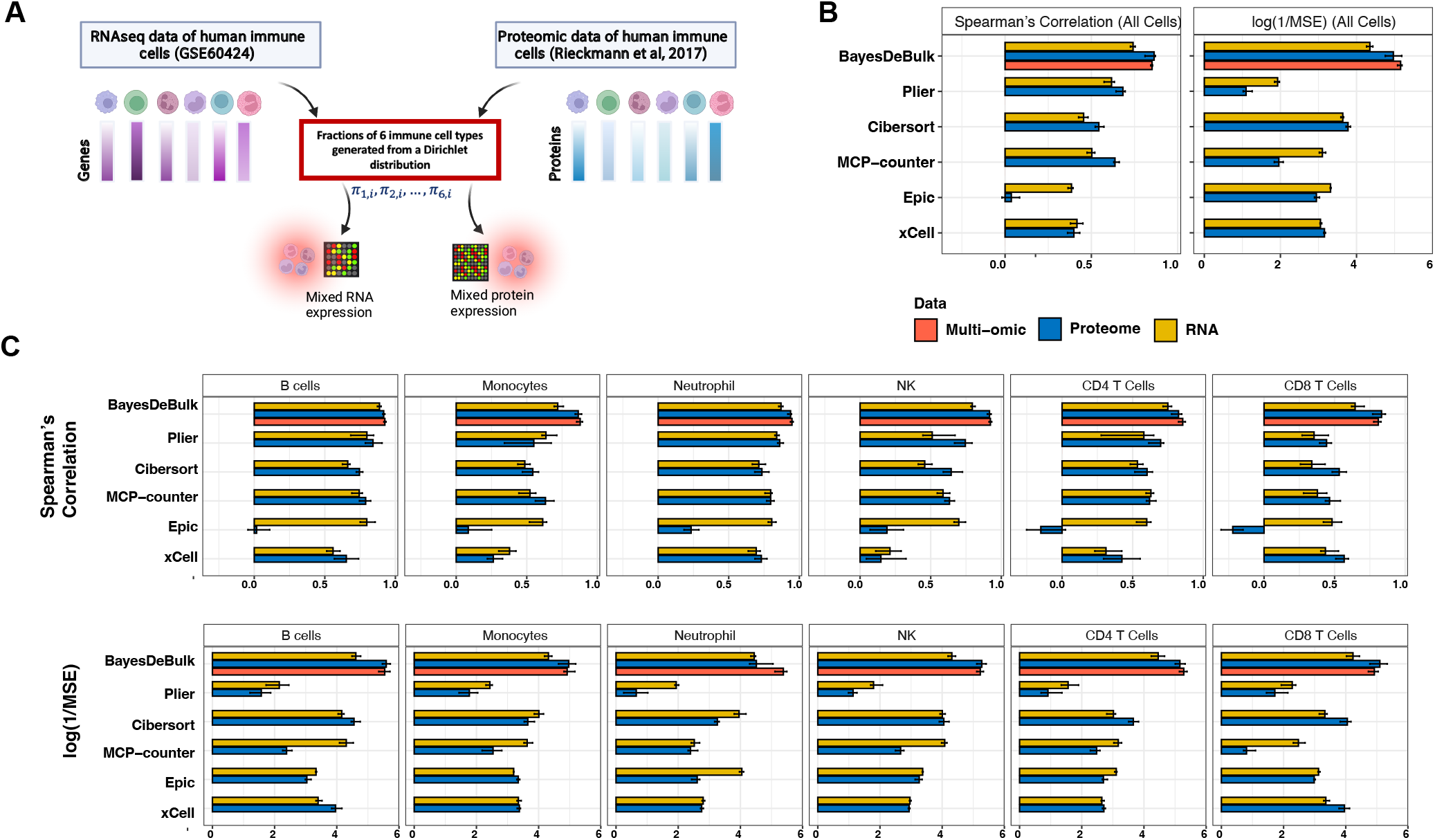
(A) Pseudo-bulk gene- and protein- expression were generated based on immune profiles from two publicly available data sets. The same set of mixture weights drawn from a Dirichlet distribution were considered in order to obtain pseudo-bulk data. (B-C) Spearman’s correlation and mean squared error (log(1/x) scale) between true and estimated cell-type fractions for BayesDeBulk, Cibersort, EPIC, MCP-counter, Plier and xCell. Barplots correspond to median values, while error bars to interquartile range across 30 replicates. All algorithms were implemented based on proteomic (Pro) and transcriptomic (RNA) data. For BayesDeBulk, a multi-omic based deconvolution was also performed (Multi-omic).

### 2.3. Real Data

#### 2.3.1. Validation based on flow cytometry

In this section, the performance of BayesDeBulk is compared with Plier [16], Cibersort [5], xCell [15], CibersortX [9], EPIC [6] and MCP-counter [10] based on transcriptomic data from peripheral blood mononuclear cells. For this comparison, we used two public gene expression data based on influenza vaccination cohorts referred to as influenza cohort 1 [30] and influenza cohort 2 [31, 32, 33]. As additional cohort, we used the gene expression of a peripheral blood data cohort involving 20 patients [5]. For BayesDeBulk, Epic, Plier and Cibersort inference the LM22 signature matrix from Cibesort was considered. For MCP-counter and xCell estimation, their default signature was utilized. Further details on how the signature matrix was leveraged for different algorithms can be found in the Star Method.

Figure 4 shows the Spearman’s correlation and MSE between flow-cytometry data and estimates derived using different algorithms. BayesDeBulk resulted in an overall Spearman’s correlation higher than all other algorithms for the influenza cohort 1. Cibersort and CibersortX models performed poorly in estimating the fraction of Monocytes. All algorithms performed well in estimating the fraction of B cells and CD8 T cells. For influenza cohort 2, BayesDeBulk was outperformed only by MCP-counter in terms of both Spearman’s correlation and mean squared error. Again, Cibersort and CibersortX performed poorly in estimating Monocytes fractions. For both data sets, MCP-counter and BayesDeBulk resulted in the lowest MSE considering all cell types combined. Finally, for the peripheral blood data involving 20 samples, BayesDeBulk was outperformed only by Cibersort and CibersortX in terms of Spearman’s correlation. Plier was not able to map any estimated factors to CD8 T cells; and therefore summary statistics for this cell type were not reported. The overall Spearman’s correlation considering the remaining cell types was lower than that of BayesDeBulk (spearman’s correlation of 0.40 versus 0.45). For all three datasets, Plier was the algorithm resulting in the worst mean squared error. Since BayesDeBulk performs the estimation of both cell-type fractions and cell-type gene expression, we expect BayesDeBulk to perform less accurately for small number of samples compared to algorithms using a fixed signature matrix (e.g., Cibersort). However, as shown by this data example, BayesDeBulk is still among the top performers in estimating cell-type fractions for datasets involving a small number of samples (n=20).

**Figure 4:**
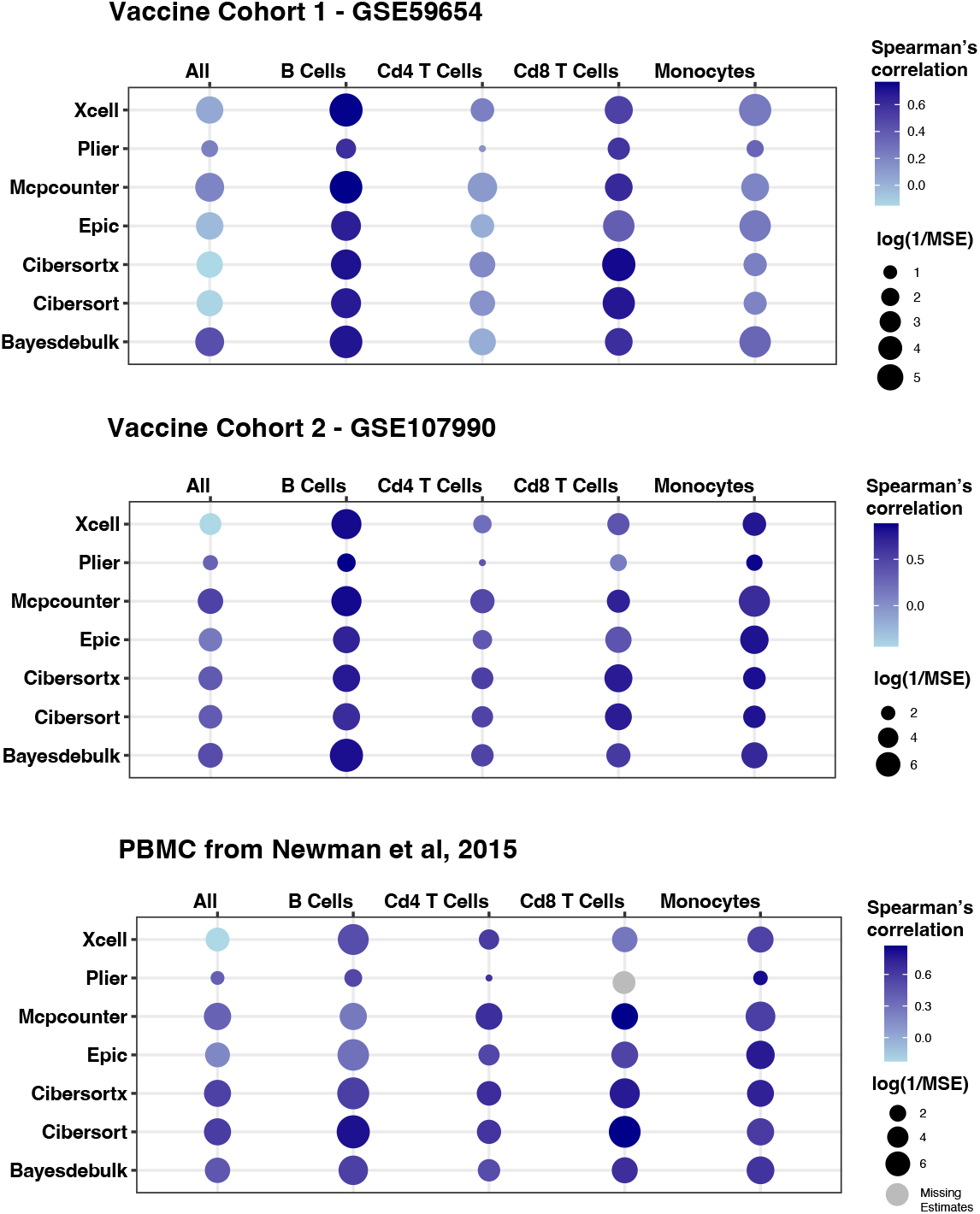
Spearman’s correlation and MSE between estimated cell-type fractions and estimates based on flow-cytometry for BayesDeBulk, Cibersort, CibersortX, Plier, EPIC and MCP-counter for different datasets

#### 2.3.2. Validation via immunohistochemistry

We considered the proteogenomic data from a recent CPTAC ovarian cancer study [34], in which RNAseq and mass spectrometry-based proteomic profiling were applied to 168 Formalin-Fixed Paraffin-Embedded (FFPE) ovarian tumor samples (*http://ptrc.cptac − data − view.org/*). While proteomics data were successful collected for all tumor samples, qualified sequencing libraries and RNAseq data were obtained for only a subset of samples (106/168). A lot of factors contribute to the challenges in performing RNAseq profiling on FFPE samples. For example, it has been reported that the time between sample acquisition and effective fixation, cellular processes and tissue autolysis can all lead to a degradation of RNA [35, 36]. Due to the sub-optimal quality of the RNAseq data, it is of interest to perform the deconvolution using the proteomic data in this study. Specifically, we applied BayesDeBulk to estimate cell-type fractions via the integration of proteomic and gene expression data, while applied other methods (xCell, Cibersort, Epic and McpCounter) to the proteomic data set instead of the RNAseq data. Note, when applying BayesDeBulk to multi-omic data, BayesDeBulk can handle omics data sets with different samples sizes (Star Methods). Such a flexibility is undoubtedly very important for real data analysis.

To validate cell type-fraction estimates, we evaluated CD8, CD4 and CCR5 expression by multiplex immunohistochemistry (IHC) for a subset of 11 samples with remaining tumor tissue blocks (Star Method). CD8 marker is expressed by CD8 T cells and can validate effectively the presence of CD8 T cell in the tumor microenvironment. On the other hand, CD4 and CCR5 are expressed in T Cells and Macrophage. As shown by Figure 5, BayesDeBulk resulted in a higher Spearman’s correlation between CD8 with CD8 T cell fraction and between CCR5 and macrophage fractions revealing the superior performance of BayesDeBulk in estimating these cell types compared to other algorithms. Cibersort was the algorithm resulting in the worst correlation between cell type fraction and corresponding cell type markers.

**Figure 5:**
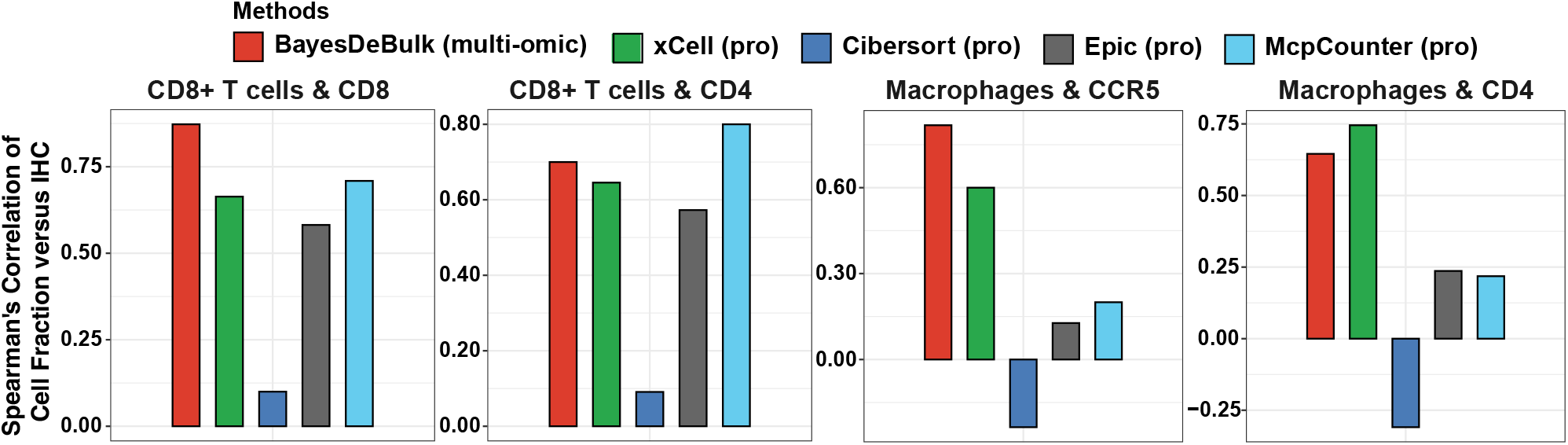
Real data validation via Immunohistochemistry. Bar-plot showing the Spearman’s correlation between cell-type fractions and marker staining via immunohistochemistry. Cell-type fractions was evaluated via BayesDeBulk (red), xCell (green), Epic (gray), Cibersort (blue) and McpCounter (cyan). For each method, correlation between CD8 T cell fractions versus CD8 and CD4 genes and between macrophages fractions versus CCR5 and CD4 genes are reported.

#### 2.3.3. Proteogenomics characterization of renal cancer

We examined proteogenomic multi-omics data from 151 kidney tumors including 103 ccRCC from Clark et al [2], 48 non-ccRCC tumors, and 101 benign adjacent kidney tissue (NAT) samples (79 from ccRCC [2] and 22 non-ccRCC patients). The 48 non-ccRCC patients represent various histologic subtypes of which 41 patients were newly profiled and seven were from our previous study [2]. For all the samples, proteomic and RNAseq data were generated.

We performed immune deconvolution analysis based on both RNA and protein data via BayesDeBulk to study the immune composition of these kidney tumors (Figure 6A). Overall, the extent of immune infiltration was lower in non-ccRCC than in ccRCC. We performed the deconvolution based on other algorithms such as EPIC, xCell and Cibersort based on gene expression data. The performance of different algorithms was assessed based on the association with progression free survival (PFS). In particular, we estimated two Cox-regression models: 1) considering only ccRCC tumors, and 2) considering all tumors types after adjusting by non-ccRCC/ccRCC status. Macrophages A cells were found significantly associated with PFS only based on BayesDeBulk multiomic deconvolution (Bonferroni’s adjusted p-values < 10%). This association with PFS is also showed by the Kaplen-Meier (KM) curve in Figure 6C. Other algorithms instead failed to capture this association which was already reported in the literature [37]. Althogh CD8 T cells did not survive the Bonferroni’s adjustment (Fig. 6B), KM curves in Figure S5 shows that BayesDeBulk-derived CD8 T cells resulted in better PFS and this association was better captured by BayesDeBulk multi-omic compared to other algorithms. These findings highlight the power of a multi-omic based deconvolution also in terms of association with patient prognosis.

**Figure 6:**
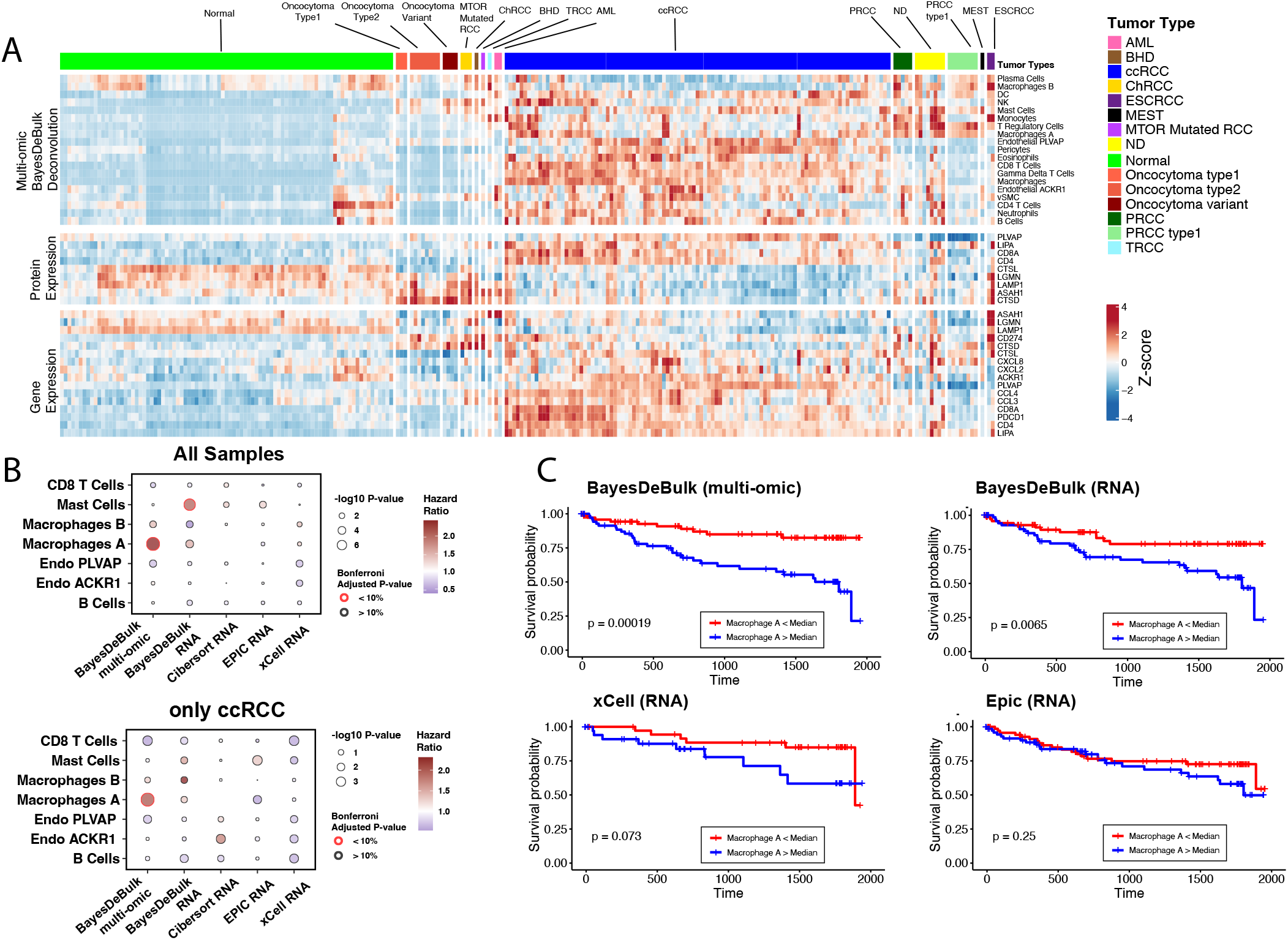
Deconvolution of Renal Cancer. A. Deconvolution of ccRCC and nonccRCC samples via the integration of proteomic and RNAseq data via BayesDeBulk. The top part of the heatmap shows the fraction of different cell-types for different samples (Multi-omic BayesDeBulk Deconvolution); while the lower part of the heatmap shows the gene expression and protein expression of some cell-type specific markers. Different tumor types are displayed in the annotation track on the top of the heatmap. B. Association between cell-type fractions and progression free survival via Cox Proportional Hazards Regression model. Cell type fractions have been estimated via BayesDeBulk integrating proteomic and RNAseq data (BayesDeBulk multi-omic), BayesDeBulk considering RNAseq data only (BayesDeBulk RNA), Cibersort, xCell and Epic based on RNAseq data (Cibersort RNA, xCell RNA, Epic RNA). The color of the bubble corresponds to the Hazard Ratio, while the size of the bubble to the p-value from the regression model (-log10 transformed). Associations with a Bonferroni adjusted p-value lower than 10% are displayed with a red outer circle. Association analysis is displayed for all samples (All Samples) and considering ccRCC samples only (only ccRCC). C. Kaplen-Meier plot showing the association between Macrophages A fraction and PFS using all renal cancer samples. Cell type fractions have been estimated via BayesDeBulk integrating proteomic and RNAseq data (BayesDeBulk multi-omic), BayesDeBulk, xCell and Epic considering RNAseq data only (BayesDeBulk RNA, xCell RNA, Epic RNA).

## Acknowledgement

This work was supported in part through the computational and data resources and staff expertise provided by Scientific Computing and Data at the Icahn School of Medicine at Mount Sinai and supported by the Clinical and Translational Science Awards (CTSA) grant ULTR004419 from the National Center for Advancing Translational Sciences.

This work was supported by grant U24CA210955, U24CA210993, U01CA214114, U24CA271114 from the National Cancer Institute Clinical Proteomic Tumor Analysis Consortium (CPTAC). Additional funding support was provided by 1F30CA265288, HHMI Gilliam Fellowship GT15758; and Associazione Italiana Ricerca sul Cancro (AIRC) under IG 2018 - ID. 21846 and AIRC 5 per Mille 2018 - ID.21073.

## Conclusion

We introduced BayesDeBulk, a novel Bayesian method for deconvolution analysis of bulk omics data. BayesDeBulk employs a mixture model to simultaneously estimate cell fractions and mean values of marker expressions of different cell types in bulk tissues. Unlike reference-based methods, BayesDeBulk does not require as input the expression levels of cell type-specific markers, making it well-suited for deconvolusion analysis of proteomic data, because properly-scaled reference protein signatures of purified cells is often difficult to obtain. Furthermore, to the best of our knowledge, BayesDeBulk is the first algorithm that enables deconvolution analysis based on multi-omics data. The advantage of jointly modeling matched RNA and proteomics data by BayesDeBulk has been demonstrated in multiple data examples.

BayesDeBulk uses a novel strategy to incorporate prior knowledge on cell type-specific marker information. Specifically, BayesDeBulk considers as input lists of differentially expressed markers between each pair of cell types (e.g., based on reference data sets of purified cells or single-cell sequencing data). Then, it imposes Repulsive priors on the mean values of these markers to ensure that the desired up/downexpression patterns are achieved between the corresponding two cell types. Compared to commonly used strategy that focuses on lists of markers that are over-expressed in one cell type compared to all others types, the approach used in BayesDeBulk is more effective to characterize subtle differences between cell types. For instance, the set of markers distinguishing CD8+ T cells from Myeloid cells would be distinct from the set that differentiates CD8+ T cells from T Regulatory cells. Additionally, the use of Repulsive prior in BayesDeBulk ensures the identifiability of the mixture model and uniquely determines the mapping between the mixture components and various cell types. In the contrast, reference-free methods based on factor analysis often suffer from identifiability issues.

The performance of BayesDeBulk was compared with that of Cibersort(X), Epic, xCell, MCP-counter and Plier using various synthetic data examples. Overall, BayesDeBulk resulted in superior performance compared to other algorithms based on pseudo-bulk data generated from Gaussian distributions as well as data based on mixtures of protein/gene expression profiles of purified cells. In particular, we demonstrated that the multi-omic deconvolution resulted in superior performance compared to single-omic deconvolution, confirming the importance of multi-omic data integration.

The superior performance of BayesDeBulk is also illustrated using three real omics datasets collected from different tissue types: transcriptomic data of peripheral blood mononuclear cells, proteogenomic data of FFPE ovarian cancer samples, and proteogenomic data of fresh frozen renal cancer samples. In the peripheral blood data example, we leveraged three public gene expression data sets based on influenza vaccination cohorts [30, 31, 32, 33], and validated cell type fractions estimates via matched flow cytometry data on a subset of the samples. BayesDeBulk was found to have on average higher correlation and lower mean squared error than other algorithms when estimating cell fractions compared to flow cytometry measurements.

The FFPE ovarian proteogenomic data example utilized IHC staining of four immune cell markers to validate the cell type fraction estimates, and demonstrated superior performance of BayesDeBulk over other algorithms. The large archives of FFPE tissue specimens offer a wealth of resources for research. Having an effective algorithm, such as BayesDeBulk, for deconvolution analysis based on FFPE-derived omics data could greatly facilitate a broad range of studies.

In the last application of renal cancer proteogenomic study, we applied BayesDeBulk and other deconvolution tools to analyze the proteogenomic data of 151 renal tumors. Notably, cell-type estimates from BayesDeBulk showed stronger associated with progression free survival in renal cancer patients than other methods. This observation highlights the importance of accurate cell-type fraction estimates in gaining insights into disease prognosis.

We have developed a user-friendly freely available web-platform to facilitate the implementation of BayesDeBulk. This web-platform will perform: (1) the estimation of the fraction of different cell types, (2) the estimation of the expression of different markers for multiple cell types. The user will be able to implement the deconvolution based on either gene expression or proteomic data and the integration of both data types. This platform is web-based for optimal portability between hardware environments, without the need for specialized or complicated installation steps from the users.

Computational time strictly depends on the number of samples and the number of cell types considered as input. The minimum number of MCMC iterations recommended is 5,000 with a burn-in of 1,000. As an example, the computational time for the deconvolution of gene expression data for 150 samples and 22 cell types implemented on Intel 8186 core at 2.7 GHz based on those parameters is around 3 hours and a half.

With the advancement of sequencing technology, it is more and more common to obtain multi-omic data for the same set of biological samples. Besides proteomic and RNAseq data, BayesDeBulk could be easily extended to analyze/integrate other data types such as methylation profiles and single cell omics data measured for the same set of tissue samples.

Since BayesDeBulk estimates cell-type fractions and mean expression levels of markers in different cell types, the number of parameters in the model increases quickly with the number of cell types. For this reason, we encourage the user to estimate a moderate number of cells (around 20), especially when working with a moderate sample size (*n <* 100).

