## Supplement Document for "BayesDeBulk: A Flexible Bayesian Algorithm for the Deconvolution of Bulk Tumor Data"

### 1. BayesDeBulk

#### *1.1. Prior distribution*

Standard choice for the parameters of the Gaussian prior is $\xi_{k,j} = 0$ and $\lambda_{k,j} = 1$; alternatively, those parameters might be chosen based on prior knowledge from existing databases and single-cell datasets. For ease of computation, we will not require $\{\pi_{i,k}\}_{k=1}^{K}$ to sum to 1. However, we will require these parameters to be defined on the unit interval $[0, 1]$. As prior specification, we will use a spike-and-slab prior [1] defined on the unit interval, i.e., $\pi_{i,k} \sim w_k N_{[0,1]}(0, 0.0001) + (1 - w_k) N_{[0,1]}(0, \gamma_k)$ with $w_k \sim Beta(1, 1)$ and $\gamma_k \sim \text{Inverse-Gamma}(a_\gamma, b_\gamma)$. The spike component concentrates its mass at values close to zero, shrinking small effects to zero and inducing sparsity in the estimates $\{\pi_{i,k}\}$. Since some cell types will be more abundant (i.e., different from zeros) than others, the percentage of zero values, i.e., $w_k$, will vary across different cell types. For instance, T cells will be more likely present in kidney or lung tissues rather than brain tissues. For the variance components $\{\sigma_j\}_{j=1}^{p}$, standard inverse-gamma priors will be utilized. Posterior computation of model parameters will be performed via Gibbs sampling [2].

### 2. Supplementary section

#### *2.1. Gibbs Sampling Steps*

Posterior computation of model parameters will be performed via Gibbs sampling [2]. Following Petralia et al [3], a latent variable $\rho$ will be introduced to facilitate the sampling from the Repulsive prior. This latent variable will be jointly modeled with $\boldsymbol{\mu}$ through the following multivariate density:

$$p(\boldsymbol{\mu}, \rho) = \left[ \prod_{\forall k; \forall j} \mathrm{N}(\mu_{k,j}; \xi_{k,j}, \lambda_{k,j}) \right] 1(h(\boldsymbol{\mu}_1, \ldots \boldsymbol{\mu}_K) > \rho)$$

A set of additional latent variables $\{Z_{i,k}\}$ will be introduced in order to facilitate the sampling from the spike-and-slab prior placed on $\{\pi_{i,k}\}$. In particular, $Z_{i,k}$ will be equal to 1 if $\pi_{i,k}$ will be sampled from the "spike" component, i.e., $\pi_{i,k} \sim N_{[0,1]}(0, 0.0001)$; while equal to 0 if $\pi_{i,k}$ will be sampled from the "slab" component, i.e., $\pi_{i,k} \sim N_{[0,1]}(0, \gamma_k)$. The Gibbs sampling scheme can be summarized in the following steps:

Step 1 Sample mean parameter $\mu_{k,j}$ from a truncated Gaussian distribution:

$$\mu_{k,j} \sim N\left( \left( \frac{\sum_{i=1}^{n} M_{i,j}\pi_{i,k}}{\sigma_j} + \frac{\xi_{k,j}}{\lambda_{k,j}} \right) \left( \frac{1}{\lambda_{k,j}} + \frac{\sum_{i=1}^{n} \pi_{i,k}^2}{\sigma_j} \right)^{-1}, \left( \frac{1}{\lambda_{k,j}} + \frac{\sum_{i=1}^{n} \pi_{i,k}^2}{\sigma_j} \right)^{-1} \right) 1(\mu_{k,j} \in S_{k,j})$$

with $M_{i,j} = y_{i,j} - \sum_{s \neq k} \mu_{s,j}\pi_{i,s}$ and $S_{k,j}$ being defined as the intersection across all constraints involving $\mu_{k,j}$. This set is defined in Section 1.3.

Step 2 Sample $Z_{i,k}$ from

$$Z_{i,k} \sim Binomial\left( \frac{w_k N_{[0,1]}(\pi_{i,k}; 0, 0.0001)}{w_k N_{[0,1]}(\pi_{i,k}; 0, 0.0001) + (1 - w_k) N_{[0,1]}(\pi_{i,k}; 0, \gamma_k)} \right)$$

Step 3 Sample $\pi_{i,k}$ from a truncated univariate Gaussian defined as:

$$N_{[0,1]}\left( \sum_{j=1}^{p} \frac{M_{i,j}\mu_{k,j}}{\sigma_j} \left( \sum_{j=1}^{p} \frac{\mu_{k,j}^2}{\sigma_j} + \frac{1}{\eta_k} \right)^{-1}, \left( \sum_{j=1}^{p} \frac{\mu_{k,j}^2}{\sigma_j} + \frac{1}{\eta_k} \right)^{-1} \right)$$

with $\eta_k = \gamma_k$ if $\ell = 0$ and $\eta_k = 0.0001$ if $\ell = 1$.

Step 4 Sample $w_k$ from

$$Beta\left(1+\sum_i 1(Z_{i,k}=1), 1+\sum_i 1(Z_{i,k}=0)\right)$$

Step 5 Sample $\gamma_k$ from:

$$\text{Inverse-Gamma}\left(\alpha_\gamma+\sum_i 1(Z_{i,k}=0)/2, \beta_\gamma+0.5\sum_{i|Z_{i,k}=0}\pi_{i,k}^2\right)$$

Step 6 Sample $\sigma_j$ from:

$$\text{Inverse-Gamma}\left(\alpha_\sigma+n/2, \beta_\sigma+0.5\sum_{i=1}^{n}\left(y_{i,j}-\sum_{k=1}^{K}\mu_{k,j}\pi_{i,k}\right)^2\right)$$

Step 7 Sample $\rho$ from a uniform distribution

$$(\rho|-)\sim Uniform(0, h(\boldsymbol{\mu}_1,\ldots,\boldsymbol{\mu}_K))$$

*2.2. Likelihood and Full Conditionals*

The likelihood of BayesDeBulk model is defined as:

$$f(\boldsymbol{Y})=\prod_{i=1}^{n}\prod_{j=1}^{p}(\sigma_j 2\pi)^{-1/2}\text{exp}\left(-\frac{1}{\sigma_j}\left(y_{i,j}-\sum_{k=1}^{K}\pi_{i,k}\mu_{k,j}\right)^2\right)$$

**Full conditional of** $\mu$ The full conditional of the mean expression of the $j$-th marker for the $k$-th cell-type (i.e., $\mu_{k,j}$) is derived as follows:

$$p(\mu_{k,j}|-)\propto \exp\left(-\sum_{i=1}^{n}\frac{1}{2\sigma_j}\left(y_{i,j}-\sum_{s=1}^{K}\mu_{s,j}\pi_{i,s}\right)^2\right)\exp\left(-\frac{1}{2\lambda_{k,j}}(\mu_{k,j}-\xi_{k,j})^2\right)1(S_{k,j})$$

$$p(\mu_{k,j}|-)\propto \exp\left(-\sum_{i=1}^{n}\frac{1}{2\sigma_j}\left(y_{i,j}-\mu_{k,j}\pi_{i,k}-\sum_{s\neq k}\mu_{s,j}\pi_{i,s}\right)^2\right)\exp\left(-\frac{1}{2\lambda_{k,j}}(\mu_{k,j}-\xi_{k,j})^2\right)$$

Let us define $M_{i,j}=y_{i,j}-\sum_{s\neq k}\mu_{s,j}\pi_{i,s}$

$$p(\mu_{k,j}|-)\propto \exp\left(-\sum_{i=1}^{n}\frac{1}{2\sigma_j}\left(M_{i,j}-\mu_{k,j}\pi_{i,k}\right)^2\right)\exp\left(-\frac{1}{2\lambda_{k,j}}(\mu_{k,j}-\xi_{k,j})^2\right)1(S_{k,j})$$

$$p(\mu_{k,j}|-) \propto \exp\left(-\frac{1}{\sigma_j}\left(\sum_{i=1}^{n} M_{i,j}^2 + \mu_{k,j}^2\sum_{i=1}^{n}\pi_{i,k}^2 - 2\mu_{k,j}\sum_{i=1}^{n} M_{i,j}\pi_{i,k}\right)\right)\exp\left(-\frac{1}{2\lambda_{k,j}}(\mu_{k,j}-\xi_{k,j})^2\right)1(S_{k,j})$$

$$p(\mu_{k,j}|-) \propto \exp\left(-\left(\mu_{k,j}^2\left(\frac{1}{\lambda_{k,j}} + \frac{\sum_{i=1}^n \pi_{i,k}^2}{\sigma_j}\right) - 2\mu_{k,j}\left(\frac{\sum_{i=1}^n M_{i,j}\pi_{i,k}}{\sigma_j} + \frac{\xi_{k,j}}{\lambda_{k,j}}\right)\right)\right)1(S_{k,j})$$

$$\mu_{k,j} \sim N\left(\left(\frac{\sum_{i=1}^n M_{i,j}\pi_{i,k}}{\sigma_j} + \frac{\xi_{k,j}}{\lambda_{k,j}}\right)\left(\frac{1}{\lambda_{k,j}} + \frac{\sum_{i=1}^n \pi_{i,k}^2}{\sigma_j}\right)^{-1}, \left(\frac{1}{\lambda_{k,j}} + \frac{\sum_{i=1}^n \pi_{i,k}^2}{\sigma_j}\right)^{-1}\right)1(S_{k,j})$$

$S_{k,j}$ is defined as the intersection across all constraints involving $\mu_{k,j}$ contained in the Repulsive function. Letting $g(z) = \exp(-\tau z^{-\eta})$, $S_{k,j}$ is defined as $S_{k,j} = \cap_{s\neq k}\{x : x > \mu_{s,j} \,\&\, g(|\mu_{s,j} - x|) > \rho\}$ for $j \in I_k$ and $S_{k,j} = \cap_{s\neq k|j\in I_s}\{x : \mu_{s,j} > x \,\&\, g(|\mu_{s,j} - x|) > \rho\}$ for $j \notin I_k$.

**Full conditional of** $\pi$ The prior distribution of $\pi_{i,k}$ is specified as follows: $\pi_{i,k} \sim w_k N_{[0,1]}(0, 0.0001) + (1-w_k)N_{[0,1]}(0,\gamma_k)$ with $\gamma_k \sim$ Inverse-Gamma$(3, 1)$. Let $Z_{i,k} = 1$ if $\pi_{i,k} \sim N_{[0,1]}(0, 0.0001)$ and $Z_{i,k} = 0$ if $\pi_{i,k} \sim N_{[0,1]}(0,\gamma_k)$; with $Z_{i,k} \sim Binomial(w_k)$. Given this prior specification, the full conditional of $\pi_{i,k}$ is defined as

$$p(\pi_{i,k}|Z_{i,k} = \ell) \propto \exp\left(-\sum_{j=1}^{p}\frac{1}{2\sigma_j}\left(y_{i,j} - \mu_{k,j}\pi_{i,k} - \sum_{s\neq k}\mu_{s,j}\pi_{i,s}\right)^2\right)\exp\left(-\frac{1}{2\eta_k}\pi_{i,k}^2\right)$$

with $\eta_k = \gamma_k$ if $\ell = 0$ and $\eta_k = 0.0001$ otherwise. Define $T_{i,k,j} = y_{i,j} - \sum_{s\neq k}\mu_{s,j}\pi_{i,s}$, then:

$$p(\pi_{i,k}|Z_{i,k} = \ell) \propto \exp\left(-\sum_{j=1}^{p}\frac{1}{2\sigma_j}\left(T_{i,k,j} - \pi_{i,k}\mu_{k,j}\right)^2\right)\exp\left(-\frac{1}{2\eta_k}\pi_{i,k}^2\right)$$

$$[\pi_{i,k}|Z_{i,k} = \ell] \sim N\left(\sum_{j=1}^{p}\frac{T_{i,k,j}\mu_{k,j}}{\sigma_j}\left(\sum_{j=1}^{p}\frac{\mu_{k,j}^2}{\sigma_j} + \frac{1}{\eta_k}\right)^{-1}, \left(\sum_{j=1}^{p}\frac{\mu_{k,j}^2}{\sigma_j} + \frac{1}{\eta_k}\right)^{-1}\right)$$

**Full conditional of w** The full conditional of $w_k$ is defined as $Beta\left(1 + \sum_i 1(Z_{i,k} = 1), 1 + \sum_i 1(Z_{i,k} = 0)\right)$

**Full conditional of Z** The full conditional of $Z_{i,k}$ is defined as:

$$Z_{i,k} \sim Binomial\left(\frac{w_k N_{[0,1]}(\pi_{i,k}; 0, 0.0001)}{w_k N_{[0,1]}(\pi_{i,k}; 0, 0.0001) + (1 - w_k) N_{[0,1]}(\pi_{i,k}; 0, \gamma_k)}\right)$$

**Full conditional of $\gamma$** The full conditional of $\gamma_w$ is defined as:

$$\text{Inverse-Gamma}\left(\alpha_\gamma + \sum_i 1(Z_{i,k} = 0)/2, \beta_\gamma + 0.5 \sum_{i|Z_{i,k}=0} \pi_{i,k}^2\right)$$

with $\alpha_\gamma = 3$ and $\beta_\gamma = 1$.

#### *2.3. Multi-omic framework*

BayesDeBulk can be utilized to perform the deconvolution by integrating gene expression and protein expression data as illustrated by Figure 1(B). In this case, each data type can be modeled via a BayesDeBulk model, with different data-specific models sharing the same set of cell type fraction parameters. Let $y_{i,j}$ and $z_{i,j}$ be the RNA and protein expression of marker $j$ for sample $i$. The proposed multi-omic framework models $y_{i,j}$ and $z_{i,j}$ as follows:

$$y_{i,j} \sim N(\theta_{i,j}^Y, \sigma_j) \ \ , \ \ \theta_{i,j}^Y = \sum_{k=1}^K \pi_{i,k} \mu_{k,j}^Y$$

$$z_{i,j} \sim N(\theta_{i,j}^Z, \iota_j) \ \ , \ \ \theta_{i,j}^Z = \sum_{k=1}^K \pi_{i,k} \mu_{k,j}^Z$$

with $K$ being the total number of cell types, $\pi_{i,k}$ being the fraction of the $k$-th cell type for sample $i$, $\mu_{k,j}^Y$ being the expression of gene $j$ for the $k$-th cell type, $\mu_{k,j}^Z$ being the expression of protein $j$ for the $k$-th cell type. It is important to notice that the two models share the same set of cell type fractions, i.e. $\{\pi_{i,k}\}$. In this case, a Repulsive prior will be placed on the mean parameters of both models, i.e., $\{\mu_{k,j}^Y\}$ and $\{\mu_{k,j}^Z\}$. Also, given its flexible framework, BayesDeBulk allows to use one data type for samples for which both data types are not observed; and integrate both data types for samples with complete measurements.

### 3. Synthetic Data

**Data Generation** The performance of BayesDeBulk was evaluated based on synthetic data generated from a Gaussian model. We considered different simulation scenarios with varying numbers of cell-types, genes and samples; i.e., $K = 10, 20$, $p = 200, 400$, $n = 50, 100$, and variance levels $\nu$ and $\sigma$. For each synthetic scenario, 30 replicate datasets were generated and the performance of different algorithms was evaluated based on two metrics: Pearson's correlation and mean squared error (MSE) between estimated fractions and true fractions. Figure S1 summarizes how the data was generated. For each cell-type, 20 cell-type-specific markers were randomly sampled from the full list of $p$ genes. Let $I_k$ be the set of cell-type specific markers for the $k$-th cell. The mean of cell-type specific markers for a particular cell-type $k$ was sampled from a Gaussian distribution with mean randomly drawn from the interval $[1, 3]$ and standard deviation 0.5; while the mean of other markers from a Gaussian distribution centered on zero and standard deviation 0.5. The fraction of different cell-types, i.e., $(\pi_{1,i}, \ldots \pi_{K,i})$, was randomly generated from a Dirichlet distribution with parameter 0.5. Given these parameters, mixed data for the $i$-th sample was generated as follows:

$$\boldsymbol{Y}_i = \pi_{1,i}\boldsymbol{V}_{1,i} + \ldots \pi_{K,i}\boldsymbol{V}_{K,i} + \boldsymbol{\epsilon}_i$$

with $\boldsymbol{\epsilon}_i \sim N(0, \nu I)$ and $\boldsymbol{V}_{k,i} \sim N(\boldsymbol{\mu}_k, \sigma I)$.

**Prior knowledge** We implemented different algorithms for different degrees of prior knowledge on cell-type-specific markers where, for each cell-type, (i) 100% of cell-type-specific markers are known a priori and (ii) 50% of cell-type specific markers are known. Specifically, under (ii), 50% of known cell-type-specific markers were randomly drawn from the original set of cell-type specific markers. Cibersort and Epic require as input the gene expression of markers for different cell types (referred to as signature matrix). For a fair comparison, a perturbed version of the original signature matrix was considered as input. This signature matrix was generated following two approaches: (i) considering the same set of cell-type specific markers, the signature matrix was regenerated; (ii) the mean of 50% of cell-type-specific markers was randomly drawn from the interval $[1, 3]$ while the expression of the remaining 50% was sampled from a Normal with mean zero and standard deviation 0.5. Basically, after perturbation, 100% of cell-specific markers will be upregulated in a particular component following (i); while only 50% following (ii). BayesDeBulk was compared with Cibersort [4], Plier [5], xCell [6] and EPIC [7] based on different simulation scenarios with varying numbers of cell-types and markers; i.e., $(K, p, n) = (10, 200, 50)$,

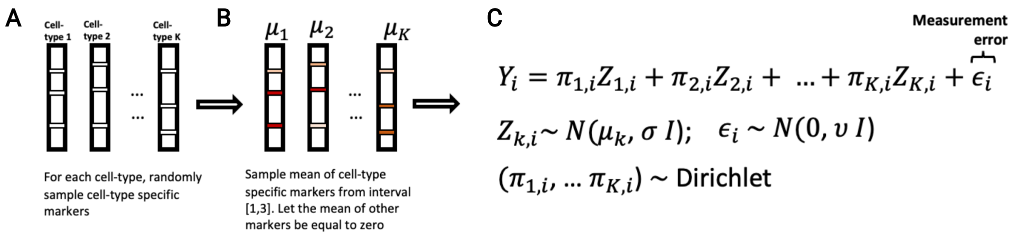


Figure S1: Synthetic data generation.

$(K, p, n) = (20, 400, 50)$, $(K, p, n) = (10, 200, 100)$ and $(K, p, n) = (20, 400, 100)$, and variance levels $\nu$ and $\sigma$. For each synthetic scenario, 30 replicate datasets were generated and the performance of different models was evaluated based on two metrics: Pearson's correlation and mean squared error (MSE) between estimated fractions and true fractions.

**Algorithms implementation** BayesDeBulk: For each synthetic data scenario, BayesDeBulk was estimated considering $10,000$ Marcov Chain Monte Carlo (MCMC) iterations; with the estimated fractions being the mean across iterations after discarding a burn-in of $1,000$. For each sample $i$, once estimated, cell-type fractions $\{\pi_{i,k}\}_{k=1}^{K}$ were standardized to sum to 1. In addition to cell-type fractions, BayesDeBulk can perform the estimation of the mean parameter for different cell types. The mean parameter $\mu_{j,k}$ was estimated for each gene $j$ and cell $k$ as the mean across 10,000 MCMC iterations after discarding a burn-in of 1,000 iterations. Plier: Plier requires as input a matrix containing cell-type specific markers. Each factor is then modeled as a function of this prior knowledge. The number of estimated factors was fixed to the total number of cell-types. One problem with Plier is that, once estimated, factors might map to multiple cell-types or to none of them. We only considered replicates for which all the factors could be uniquely mapped to a cell-type. The median number of replicates satisfying this requirement across all synthetic scenarios was 23 out of 30 with an interquartile range of $(15, 27)$. xCell xCell was implemented using the Bioconductor package GSVA [8]. Cibersort For each synthetic data, Cibersort was implemented using $P = 100$ permutations and the relative mode. Quantile normalization was not performed.

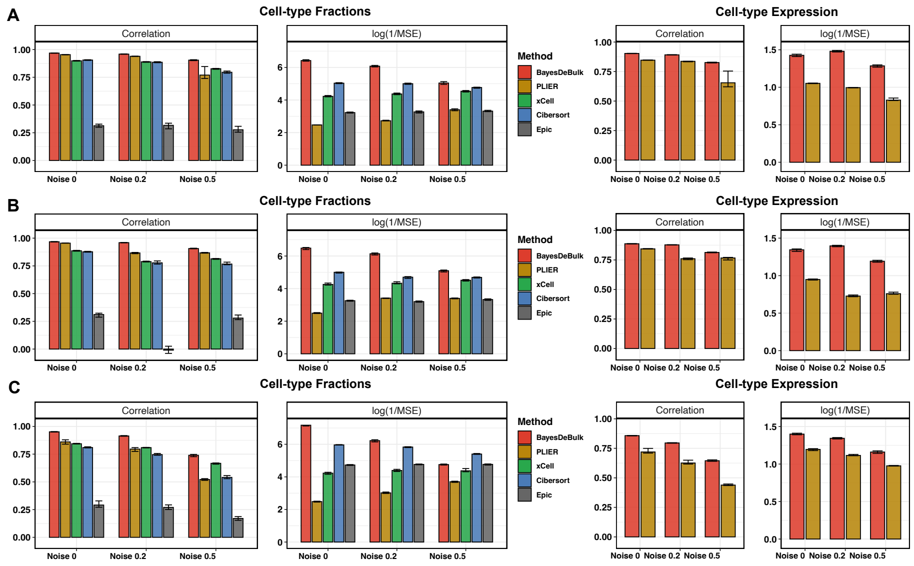


Figure S2: **Synthetic data involving 100 samples under the assumption 100% of cell-type specific markers are known** Pearson's correlation and mean squared error (MSE) between estimated values and truth over 30 replicates for BayesDeBulk (red), Cibersort (blue), xCell (green), Epic (gray) and Plier (gold). Barplots correspond to the median across different replicates while error bars to the interquartile range. For each simulation scenario, we report the correlation and MSE between the estimated cell-type fractions and truth (left-hand panel) for all five algorithms, and between the estimated cell-type expression and the truth (right-hand panel) for BayesDeBulk and Plier. Results are based on data simulated for (A) $K = 10$ and $\sigma = 0.5$; (B) $K = 10$ and $\sigma = 1$; (C) $K = 20$ and $\sigma = 0.5$ for different level of measurement errors $\nu$ (noise).

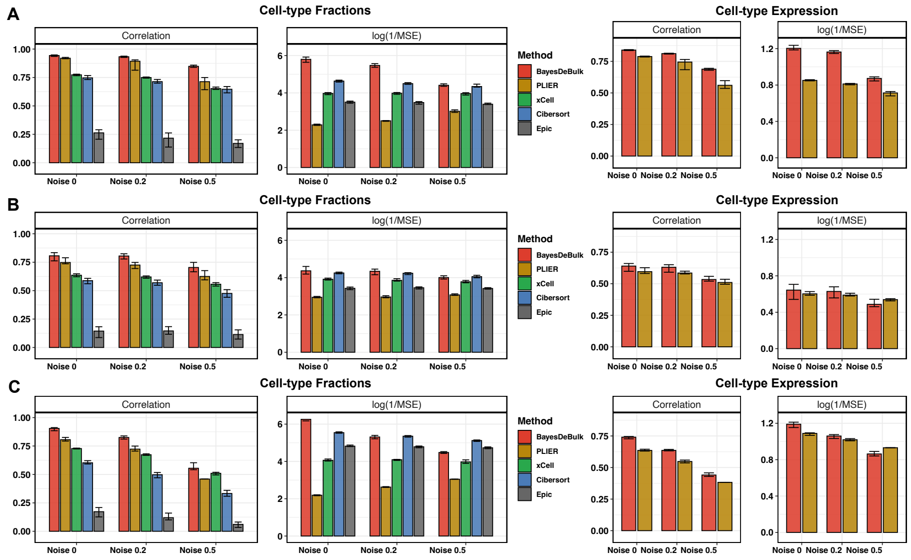


Figure S3: **Synthetic data involving 50 samples under the assumption 50% of cell-type specific markers are known** Pearson's correlation and mean squared error (MSE) between estimated values and truth over 30 replicates for BayesDeBulk (red), Cibersort (blue), xCell (green), Epic (gray) and Plier (gold). Barplots correspond to the median across different replicates while error bars to the interquartile range. For each simulation scenario, we report the correlation and MSE between the estimated cell-type fractions and truth (left-hand panel) for all five algorithms, and between the estimated cell-type expression and the truth (right-hand panel) for BayesDeBulk and Plier. Results are based on data simulated for (A) $K = 10$ and $\sigma = 0.5$; (B) $K = 10$ and $\sigma = 1$; (C) $K = 20$ and $\sigma = 0.5$ for different level of measurement errors $\nu$ (noise).

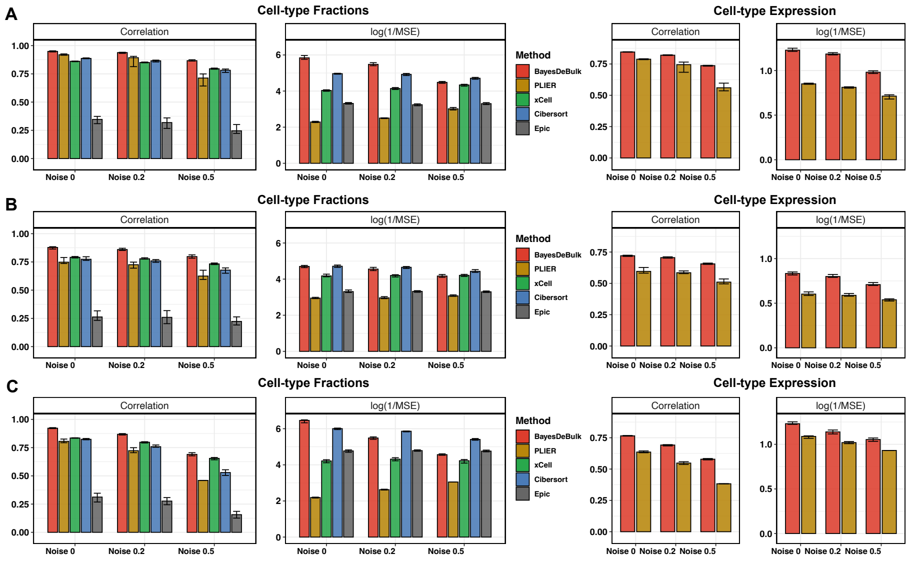


Figure S4: **Synthetic data involving 50 samples under the assumption 100% of cell-type specific markers are known** Pearson's correlation and mean squared error (MSE) between estimated values and truth over 30 replicates for BayesDeBulk (red), Cibersort (blue), xCell (green), Epic (gray) and Plier (gold). Barplots correspond to the median across different replicates while error bars to the interquartile range. For each simulation scenario, we report the correlation and MSE between the estimated cell-type fractions and truth (left-hand panel) for all five algorithms, and between the estimated cell-type expression and the truth (right-hand panel) for BayesDeBulk and Plier. Results are based on data simulated for (A) $K = 10$ and $\sigma = 0.5$; (B) $K = 10$ and $\sigma = 1$; (C) $K = 20$ and $\sigma = 0.5$ for different level of measurement errors $\nu$ (noise).

### 4. Mixture of protein/gene expression from purified cells

**Mixture of RNA expression from pure cells** We considered data from [9] which contains transcriptomic profile of $K = 6$ immune cell types such as Neutrophil, Natural Killers, B cells, CD4 T cells, CD8 T cells and Monocytes. Let $\mu_k$ be the averaged transcriptomic data across multiple replicates for the $k$-th cell type. For each sample $n$, weights of different immune cells were randomly sampled from a Dirichlet distribution with parameter 0.5 (i.e., $\pi_{n,1}, \pi_{n,2}...\pi_{n,K}$). Count data was first log2 transformed and then mixed data was derived as the weighted average of transcriptomic profile of different cell-types as follows $y_n = \sum_{k=1}^{K} \pi_{n,k} Z_{n,k} + \epsilon_n$ with $Z_{n,k} \sim N(\mu_k, \sigma)$ with $\sigma = sd(\mu)/2$ and $\epsilon_n \sim N(0, \delta)$, and $\delta$ being chosen to ensure a 1:1 signal to noise ratio.

**Mixture of protein expression from pure cells** We considered data from Rieckmann et al [10] including proteomic profile of the same set of immune cells. Considering the same set of weights $\{\pi_{n,k}\}$, mixed proteomic data was generated in a similar fashion as the transcriptomic profile.

**Prior knowledge** For the implementation of BayesDeBulk, Epic [7] , Plier [5] and Cibersort [4], the LM22 signature matrix from Cibersort was considered. For MCP-counter [11] and xCell [6] estimation, their default signatures were utilized. Contrary to other algorithms, BayesDeBulk can take as input the list of markers upregulated in one cell-type compared to another cell-type. This is a more flexible strategy than requiring a marker to be higher expressed in one cell-type compared to all other cell-types. For each pair of cells $(k, s)$, we considered a marker $\ell$ upregulated in the $k$-th cell type compared to the $s$-th cell type if $(L_k^\ell > 5 \times L_s^\ell)$ and $L_k^\ell > 1000$ with $L_k^\ell$ being the value in the LM22 signature matrix of the $\ell$ marker for the $k$-th cell type. On the other hand, Plier requires a list of markers expressed in each cell-type. For its implementation, a marker $\ell$ was considered expressed in the $k$-th cell-type if $L_k^\ell > 1000$. For the implementation of EPIC and Cibersort, the originial LM22 signature matrix was considered as input.

### 5. Validation based on cytometry immunoprofile

**Prior knowledge** For the implementation of BayesDeBulk, Epic [7] , Plier [5] and Cibersort [4], the LM22 signature matrix from Cibersort was considered [4]. For MCP-counter [11] and xCell [6] estimation, their default signatures were utilized. Contrary to other algorithms, BayesDeBulk can take as input the list of markers upregulated in one cell-type compared to another cell-type. This is a more flexible strategy than requiring a marker to be higher expressed in one cell-type compared to all other cell-types. For each pair of cells $(k, s)$, we considered a marker $\ell$ upregulated in the $k$-th cell type compared to the $s$-th cell type if $(L_k^\ell > 5 \times L_s^\ell)$ and $L_k^\ell > 1000$ with $L_k^\ell$ being the value in the LM22 signature matrix of the $\ell$ marker for the $k$-th cell type. On the other hand, Plier requires a list of markers expressed in each cell-type. For its implementation, a marker $\ell$ was considered expressed in the $k$-th cell-type if $L_k^\ell > 1000$. For the implementation of EPIC and Cibersort, the originial LM22 signature matrix was considered as input.

**Estimation** For the implementation of BayesDeBulk and Plier, the data was first log transformed and each gene was standardized to z-score (mean zero and standard deviation 1). For BayesDeBulk deconvolution, the LM22 signature matrix values were considered as prior mean $\{\xi_{k,j}\}$; while $\lambda_{k,j}$ was set to 1. BayesDeBulk model was estimated considering $10,000$ MCMC iterations; with the estimated fractions derived as the mean across iterations after discarding a burn-in of $1,000$. Once estimated, parameters $\{\pi_{i,k}\}_{k=1}^K$ were standardized to sum to 1 for each sample $i$. For CibersortX based deconvolution, a batch correction step was implemented (B-batch mode). The relative mode was utilized for both Cibersort and CibersortX deconvolutions.

### 6. Deconvolution of proteogenomic data from FFPE

*Bulk deconvolution via BayesDeBulk* We performed a multi-omic based deconvolution of bulk FFPE ovarian tumors [12] based on the integration of proteomic and RNAseq data via BayesDeBulk. For this deconvolution, we considered immune, endothelial, adipose, epithelial and fibroblast cells. In order to estimate the fraction of endothelial, epithelial, adipose and fibroblast cells, we used the gene signature derived from a single cell study [13]; while the LM22 signature matrix from Cibersort was considered in order to derive the list of cell-specific markers of immune cells [14]. For BayesDeBulk estimation, 5000 Markov-Chain Monte Carlo (MCMC) iterations were considered. The estimated fractions were derived as the median of the MCMC iterations after discarding a burn-in of 1,000 iterations. BayesDeBulk leveraged both proteomic and RNAseq data to infer cell type fractions for samples for which the two data types were available; and a single-omic deconvolution for sample with one data type observed.

*Bulk deconvolution via other algorithms* BayesDeBulk performance was compared to EPIC [7], xCell [6], Cibersort [14] and MCPcounter [11]. Similarly to BayesDeBulk, for xCell deconvolution, we utilized cell-type markers from [13] for endothelial, epithelial, adipose and fibroblast cells; and the LM22 signature matrix to derive markers for immune cells. Cell type estimates were computed via the package GSVA [8]. Epic, Cibersort and MCPcounter requires as input marker expression for different cell types. For Epic and Cibersort, the LM22 matrix was utilized [14]; while for MCPcounter the built-in signature was utilized [11]. We tried to implement PLIER [5] using the gene list used in xCell inference. However, once factors were estimated they were not uniquely mapped to different cell-types and we, therefore, decided to not report those results. Since those algorithms can only achieve a single-omic deconvolution, proteomic data, data type available for all the samples, was utilized to estimate the fraction of different cell types.

*Immunohistochemistry of ovarian tumor samples* Slide-mounted FFPE sections were dewaxed and stained on a Leica BOND Rx autostainer (Leica, Buffalo Grove, IL) using Leica Bond reagents for dewaxing (Dewax Solution), antigen retrieval and antibody stripping (Epitope Retrieval Solution 2), and rinsing after each step (Bond Wash Solution). A high stringency wash was performed after the secondary and tertiary applications using high-salt TBST solution (0.05M Tris, 0.3M NaCl, and 0.1% Tween-20, pH 7.2-7.6).

### 7. Deconvolution of renal tumors

*Deconvolution via BayesDeBulk* To estimate the fraction of different cell types in the tissue microenvironment, we performed a multi-omic based deconvolution integrating global proteomic and RNAseq data. To perform the deconvolution, BayesDeBulk requires a list of cell-type specific markers for each cell type. For immune cells, such list was derived from the LM22 signature matrix. For this analysis, an aggregated version of the LM22 signature matrix was utilized. Specifically, we averaged the LM22 values mapping to different types of CD4 T Cells (e.g., Memory T Cells, Naïve T Cells) to create a gene signature for CD4 T Cells. The same strategy was utilized for Dendritic cells, Natural Killers cells, Mast Cells and B Cells. For each pair of cell types, we considered a marker to be upregulated in the first cell type compared to the other cell type, if the corresponding value of the LM22 matrix for the first cell type was greater than 1,000 and 5 times the value of the other cell type. For Endothelial-PLVAP, Endothelial-ACKR1, Pericytes, vSMC and Tumor cells, we used marker signatures from a previous ccRCC single-cell RNAseq study [15]. To derive these signatures, differential expression between different single-cell clusters was performed and only markers significant at 10% FDR and a log fold change greater than 1 were considered as cell-type specific markers. Finally, we considered Macrophage A and Macrophage B signatures from Zhang et al [15]. As common markers for Macrophage A and B we used C1QA, C1QB, C1QC, MS4A6A, LYZ, TYROBP, FCGR2A, FCER1G, AIF1, CD14, CD68; as markers specific of Macrophages A we considered: CXCL8, CXCL2, CCL4, CCL3, CCL4L2, CXCL3, CCL3L3, CCL20, NFKB1, IL1B; while for Macrophage B the following cell-type specific markers were considered: CTSL, LGMN, ASAH1, LIPA, CTSD, LAMP1. BayesDeBulk was estimated via 10,000 Monte Carlo Markov Chain (MCMC) iterations. Cell-type fractions were estimated as the mean across MCMC iterations after discarding a burn-in of 5,000 iterations. Once estimated, cell-type fractions for each patient were standardized to sum to 1.

*Deconvolution via other algorithms* BayesDeBulk performance was compared to EPIC [7], xCell [6] and Cibersort [14]. Similarly to BayesDeBulk, for xCell deconvolution, we utilized cell-type markers from [15] for Endothelial-PLVAP, Endothelial-ACKR1, Pericytes, vSMC and Tumor cells. As common markers for Macrophage A and B we used C1QA, C1QB, C1QC, MS4A6A, LYZ, TYROBP, FCGR2A, FCER1G, AIF1, CD14, CD68; as markers specific of Macrophages A we considered: CXCL8, CXCL2, CCL4, CCL3, CCL4L2, CXCL3, CCL3L3, CCL20, NFKB1, IL1B; while for Macrophage B the following cell-type specific markers were considered: CTSL, LGMN, ASAH1, LIPA, CTSD, LAMP1. The LM22 signature matrix was instead utilized to derive markers for immune cells. In particular, for each cell type, we considered cell-type markers genes with a LM22 score greater than 1,000. Cell type estimates were

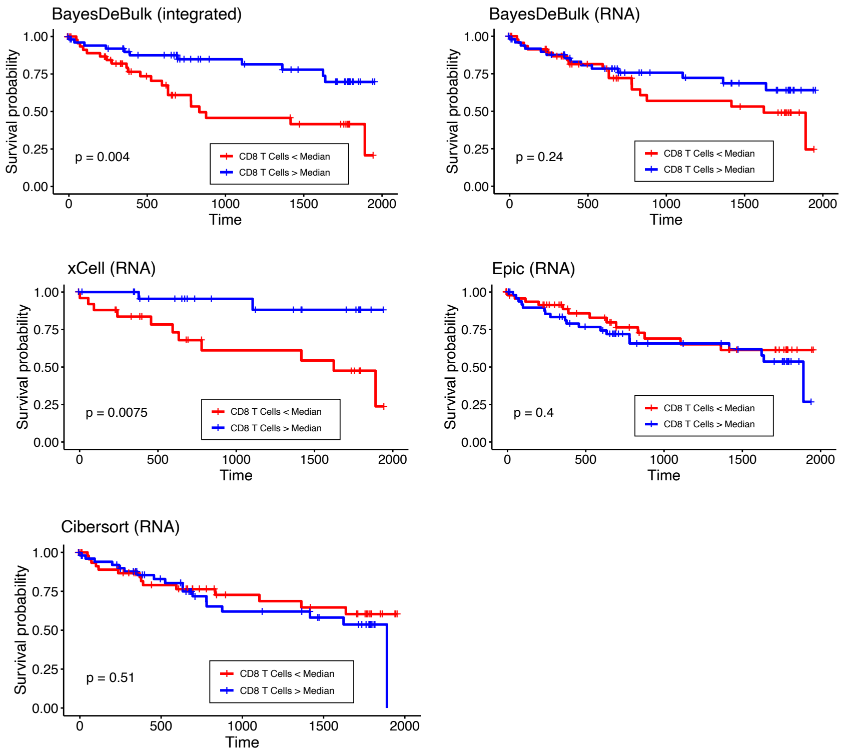


Figure S5: Kaplen-Meier plot showing the association between CD8 T cells fraction and progression free survival using ccRCC samples. Cell type fractions have been estimated via BayesDeBulk integrating global proteomic and RNAseq data (BayesDeBulk multi-omic), BayesDeBulk, xCell, Cibersort and Epic considering RNAseq data only (i.e., BayesDeBulk RNA, xCell RNA, Epic RNA)

computed via the package GSVA [8]. Epic and Cibersort requires as input marker expression for different cell types. For Epic and Cibersort, we utilized gene expression profiles for different cell types from [15]. We tried to implement PLIER [5] using the gene list used in xCell inference. However, once factors were estimated they were not uniquely mapped to different cell-types and we, therefore, decided to not report those results. Since those algorithms can only achieve a single-omic deconvolution, RNAseq data was utilized to estimate the fraction of different cell types.
